# Conserved cell-type specific signature of resilience to Alzheimer’s disease nominates role for excitatory intratelencephalic cortical neurons

**DOI:** 10.1101/2022.04.12.487877

**Authors:** Maria A. Telpoukhovskaia, Niran Hadad, Brianna Gurdon, Yanchao Dai, Andrew R. Ouellette, Sarah M. Neuner, Amy R. Dunn, Jon A. L. Willcox, Yiyang Wu, Logan Dumitrescu, Orhan Bellur, Ji-Gang Zhang, Kristen M.S. O’Connell, Eric B. Dammer, Nicholas T. Seyfried, Sukalp Muzumdar, Jesse Gillis, Paul Robson, Matthias Arnold, Timothy J. Hohman, Vivek M. Philip, Vilas Menon, Catherine C. Kaczorowski

## Abstract

Alzheimer’s disease (AD), the leading cause of dementia, affects millions of people worldwide. With no disease-modifying medication currently available, the human toll and economic costs are rising rapidly. Under current standards, a patient is diagnosed with AD when both cognitive decline and pathology (amyloid plaques and neurofibrillary tangles) are present. Remarkably, some individuals who have AD pathology remain cognitively normal. Uncovering factors that lead to “cognitive resilience” to AD is a promising path to create new targets for therapies. However, technical challenges discovering novel human resilience factors limit testing, validation, and nomination of novel drugs for AD. In this study, we use single-nucleus transcriptional profiles of postmortem cortex from human individuals with high AD pathology who were either cognitively normal (resilient) or cognitively impaired (susceptible) at time of death, as well as mouse strains that parallel these differences in cognition with high amyloid load. Our cross-species discovery approach highlights a novel role for excitatory layer 4/5 cortical neurons in promoting cognitive resilience to AD, and nominates several resilience genes that include *ATP1A1*, *GRIA3*, *KCNMA1*, and *STXBP1*. This putative cell type has been implicated in resilience in previous studies on bulk RNA-seq tissue, but our single-nucleus and cross-species approach identifies particular resilience-associated gene signatures in these cells. These novel resilience candidate genes were tested for replication in orthogonal data sets and confirmed to be correlated with cognitive resilience. Based on these gene signatures, we identified several potential mechanisms of resilience, including regulation of synaptic plasticity, axonal and dendritic development, and neurite vesicle transport along microtubules that are potentially targetable by available therapeutics. Because our discovery of resilience-associated genes in layer 4/5 cortical neurons originates from an integrated human and mouse transcriptomic space from susceptible and resilient individuals, we are positioned to test causality and perform mechanistic, validation, and pre-clinical studies in our human-relevant AD-BXD mouse panel.

## Introduction

Resilience to cognitive decline associated with Alzheimer’s disease (AD) is a phenomenon by which some people retain better than expected cognitive ability despite high amyloid and tau burdens.^1^ Bolstering the brain’s ability to cope with AD pathology is an attractive new avenue for therapies. Resilience to AD has been reported in longitudinal cohort studies, such as in the Religious Orders Study (ROS) and Memory and Aging Project (MAP) groups, in which a third of individuals with normal cognition have AD pathology.^2, 3^ While some resilience factors have been identified by studying these individuals,^4–6^ many resilience factors and the mechanisms by which they act remain unknown or poorly understood, particularly in the context of specific cell types and their contribution to resilience. In particular, a recent study^7^ used deconvolution approaches on post-mortem human bulk RNA-seq data from ROSMAP to implicate a specific neuronal subtype associated with residual cognition. Here, we extend this type of comparison by directly investigating single-nucleus RNA-seq data from human and mouse brain tissue to identify not only cell types associated with resilience, but also specific gene signature changes within these cell types. The cross-species comparison allows us to link findings to translationally relevant model organisms, which are essential to validate and test resilience candidates; such genetically diverse mouse models have already been valuable for modeling complex diseases.^8, 9^

The AD-BXDs are a panel of mice that incorporates the 5XFAD mutation into the genetically diverse BXD genetic reference panel. This panel models individual differences in memory function in response to human FAD mutations, resulting in a genetically diverse population of AD mice that is sensitive to additive effects of inherited risk loci defined by LOAD GWAS.^10^ The AD-BXD panel has been used to nominate several resilience factors based on ‘omics’ analysis of strains stratified as resilient or susceptible to cognitive decline in the presence of pathology.^11, 12^ While progress has been made using bulk data from these models (as well as human bulk RNA-seq data), there is still a need to identify and prioritize additional human-relevant resilience factors to advance drug interventions to pre-clinical studies. By harmonizing molecular signatures of resilience in this mouse model panel with those found in humans, we establish a framework to nominate translationally-relevant resilience candidates that can undergo further testing in pre-clinical studies.

In this study, we integrated cortical transcriptomic data from the human ROSMAP and the mouse AD-BXD cohorts and discovered that cognitive resilience to AD is associated with the upregulation of gene expression in a subset of excitatory neurons. We identified biological processes such as axonal and dendritic development, regulation synaptic transmission and plasticity, and axo-dendritic transport. We further narrowed down the list of candidate genes via validation in orthogonal human data sets and nominated resilience targets, including *ATP1A1*, *GRIA3*, *KCNMA1*, and *STXBP1* based on their high potential for drug targetability. Thus, we demonstrate the power of cross-species transcriptomic analyses to identify novel AD resilience factors.

## Results

### High alignment of integrated cross-species gene expression profiles enables conserved resilience factor interrogation

We integrated single nucleus transcriptomic data from human and mouse cohorts to identify conserved cell type-specific signatures of resilience to AD. We used human prefrontal cortex (PFC) tissue samples from the ROSMAP cohort,^13^ which includes tissue from a number of resilient and susceptible individuals, as well as cognitively normal controls (Fig. 1A). For mouse, we used frontal cortex tissue samples containing PFC from AD-BXD strains determined to be cognitively resilient and susceptible to the presence of the 5XFAD transgene (Fig. 1B), along with their non-transgenic genetically identical counterparts (Fig. 1A). To better align the mouse population to the human, we chose the 14-month time point and considered contextual fear memory.

**Fig. 1.**
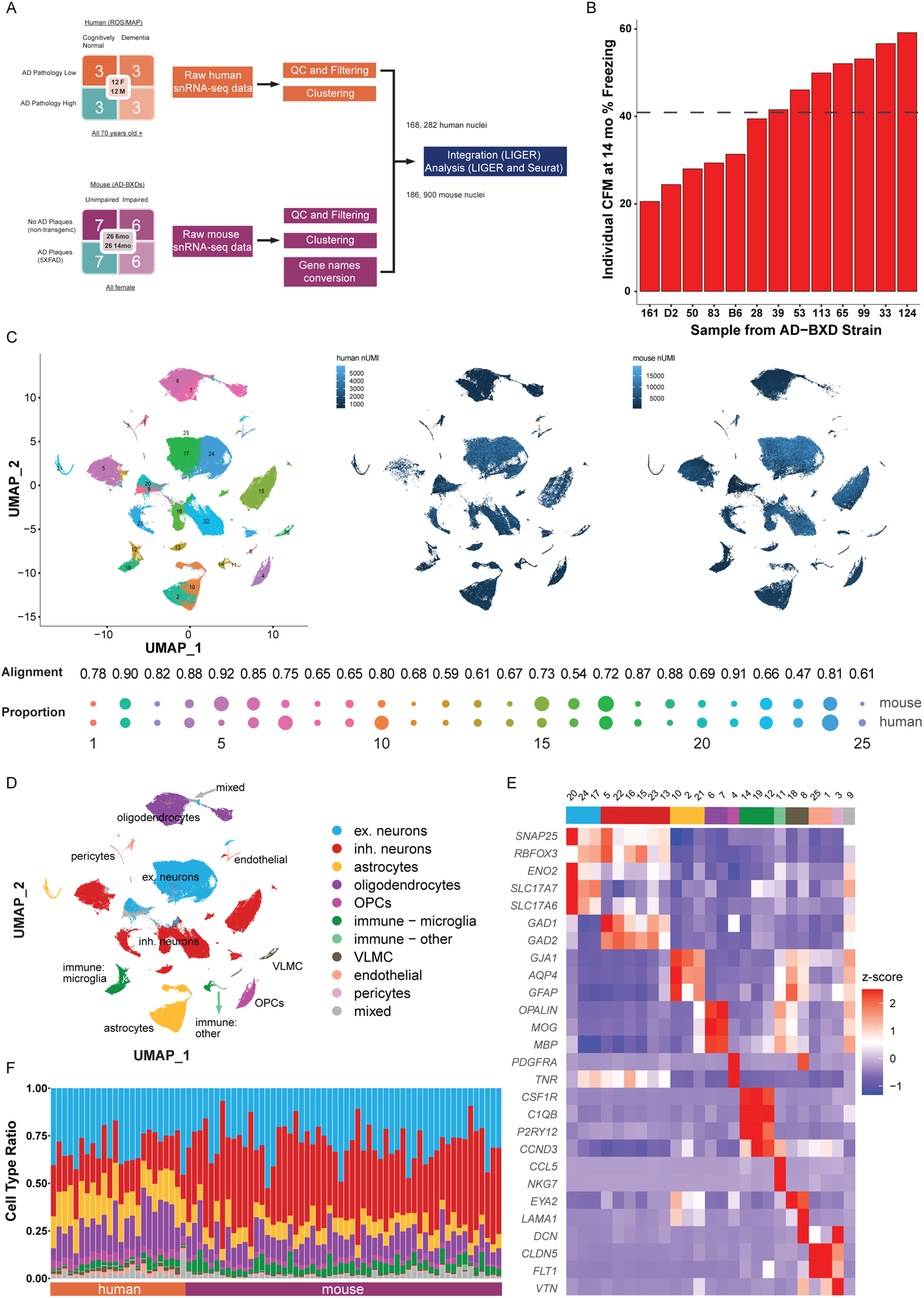
Integration of human and mouse snRNA-seq PFC data resulted in a shared data space with no compositional differences between resilient and susceptible individuals. **A.** Cohort demographics and analysis workflow for cross-species data integration; **B.** Plot of CFM values to define resilient and susceptible status for the 14 mo mice with 5XFAD transgene; **C.** Integrated human and mouse data set: UMAP visualization of the integrated data set with cluster numbers, and separate UMAP of the data for human (left) and mouse (right); alignment per cluster and cluster proportion by species below (see also Fig. S1); **D.** MetaNeighbor analysis and differential gene expression led to classification of each cluster as a cell type. Heatmap with cell-type specific marker genes demonstrate cell-type assignments. Heatmap was produced using average expression for each cluster, by default, with scaled data and maximum display value set at 2.5; **E.** UMAP for the integrated data set with cluster identity; **F.** Proportions of cell types in each sample (see also Fig. S2A).

Integration of processed data (Fig. 1A) resulted in a space with a high overall alignment score of 0.76 (comparable to examples in Welch et al 2019^14^), with individual cluster alignments that ranged from 0.47 to 0.92, with mean of 0.74 (Table S1 and Fig. 1C). The majority of clusters had alignment above 0.65, with exceptions of clusters C9 – mixed, C13 – inh. neurons, C25 – endothelial, C12 - immune – microglia, C16 – inh. neurons, and C23 – inh. neurons, which had lower confidence scores. The lower alignment scores for inhibitory neurons may be indicative of tissue dissection in mouse including subcortical regions. There was also comparable representation of human and mouse nuclei in 25 clusters (as observed in cluster proportion composition and individual UMAPs in Fig. 1C). Additionally, all clusters had contributions from human and mouse samples (Fig. S2A).

All major cell types were identified in the integrated data set using MetaNeighbor and marker genes: excitatory (clusters 17, 20, and 24) and inhibitory (clusters 5, 13, 15, 16, 22, and 23) neurons, astrocytes (cluster 2, 10, and 21), oligodendrocytes (clusters 6 and 7), oligodendrocyte precursors (OPCs, cluster 4), microglia (clusters 12, 14, and 19), vascular and leptomeningeal cells (VLMC, clusters 8 and 18), endothelial cells (clusters 1 and 25), and pericytes (cluster 3) (Fig. 1D and E, Table S2). Cluster 9 had a mixed cell signature and was labeled as a “mixed cluster” (Fig. 1E, rightmost cluster). The ratio of cell types across samples was proportionally similar (Fig. 1F). Additionally, we observed high correlation values within each cluster for human vs mouse when gene expression in each cluster was correlated between species (Fig. 2A and B, top and center), with lower correlation between clusters assigned to different cell types (Fig. 2A and B, bottom). We also confirmed cell type cluster assignment using marker genes when the data were separated by species (Fig. 2C). Note that because the mouse gene names were converted to their human equivalent, when describing the mouse portion of results from the integrated data analysis, the names were left in all capital letters. When the mouse results were considered separately, homologous mouse gene names were used.

**Fig. 2.**
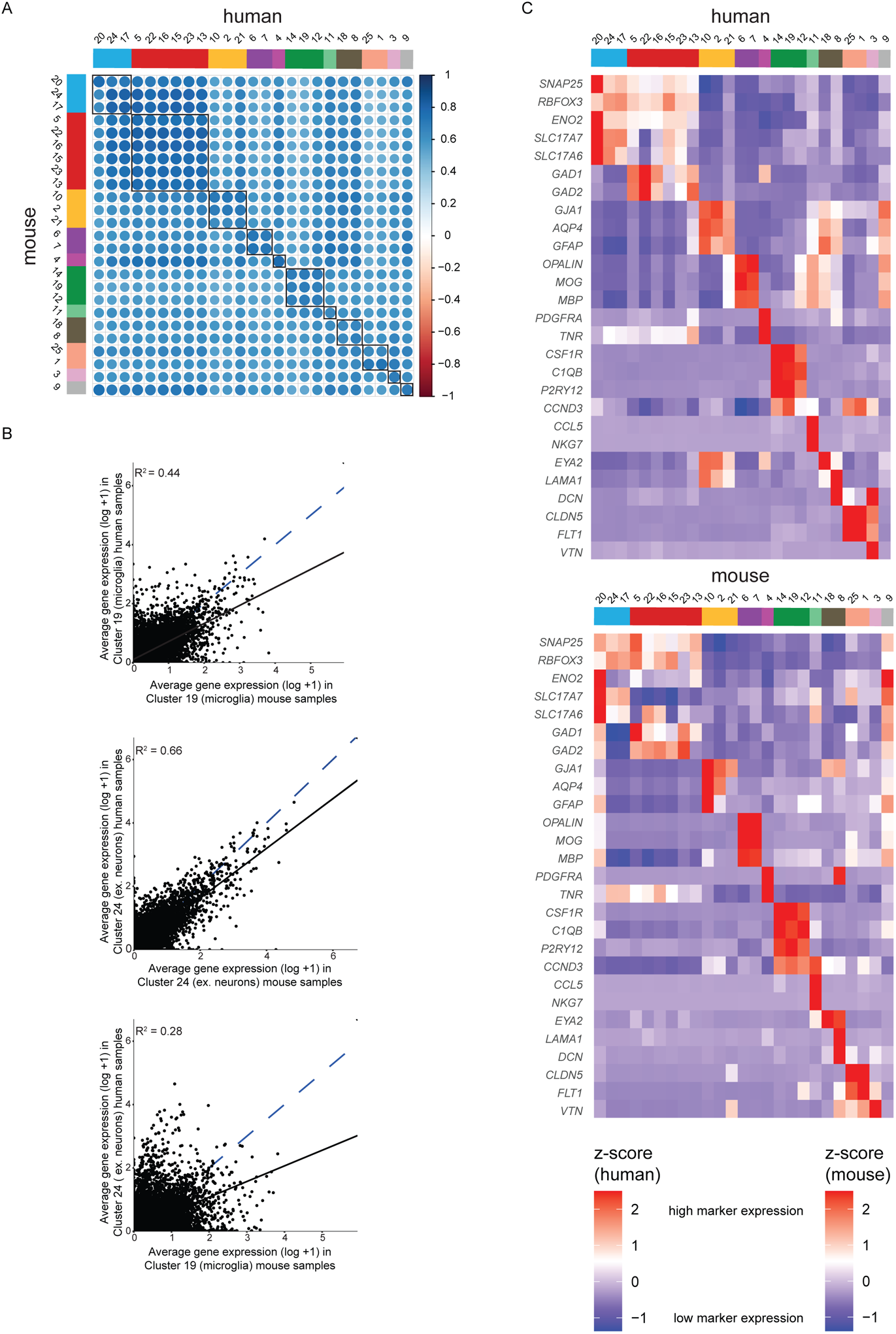
Correlation analysis of genes per cluster and separate UMAPs for each species. **A.** Correlation plot of genes in each cluster across species; **B.** Example correlations showing each gene for a microglia cluster in human and mouse, for an ex. neuronal cluster for human and mouse, and for a microglia mouse cluster vs a human neuronal cluster with lower correlation value, with dashed blue line representing perfect concordance (slope of 1) and the solid black line the fit of the data used to calculate R^2^ value; **C.** Individual heatmaps for human and mouse data within the integrated clusters demonstrating a shared class identity in all but one cluster (right most cluster, #9).

### Gene upregulation in excitatory neurons is the strongest resilience signature

Resilience may arise due to broad changes in cellular composition, changes in gene expression, or a combination of both. Thus, to probe the nature of resilience in the AD-BXD population, we analyzed whether resilience is associated with differences in cellular composition and/or gene expression. We found that there were no statistically significant differences in cluster composition between resilient and susceptible individuals in the human cohort and in the 14-month-old AD-BXDs (Fig. 3A and B). This suggests that changes associated with resilience are subtle (at the resolution of our clustering), leading us to hypothesize that the differences in long term memory between resilient and susceptible individuals may be reflected in altered gene expression in one or more cell types, and/or changes in the proportion of nuclei expressing a set of genes within one or more cell types.

**Fig. 3.**
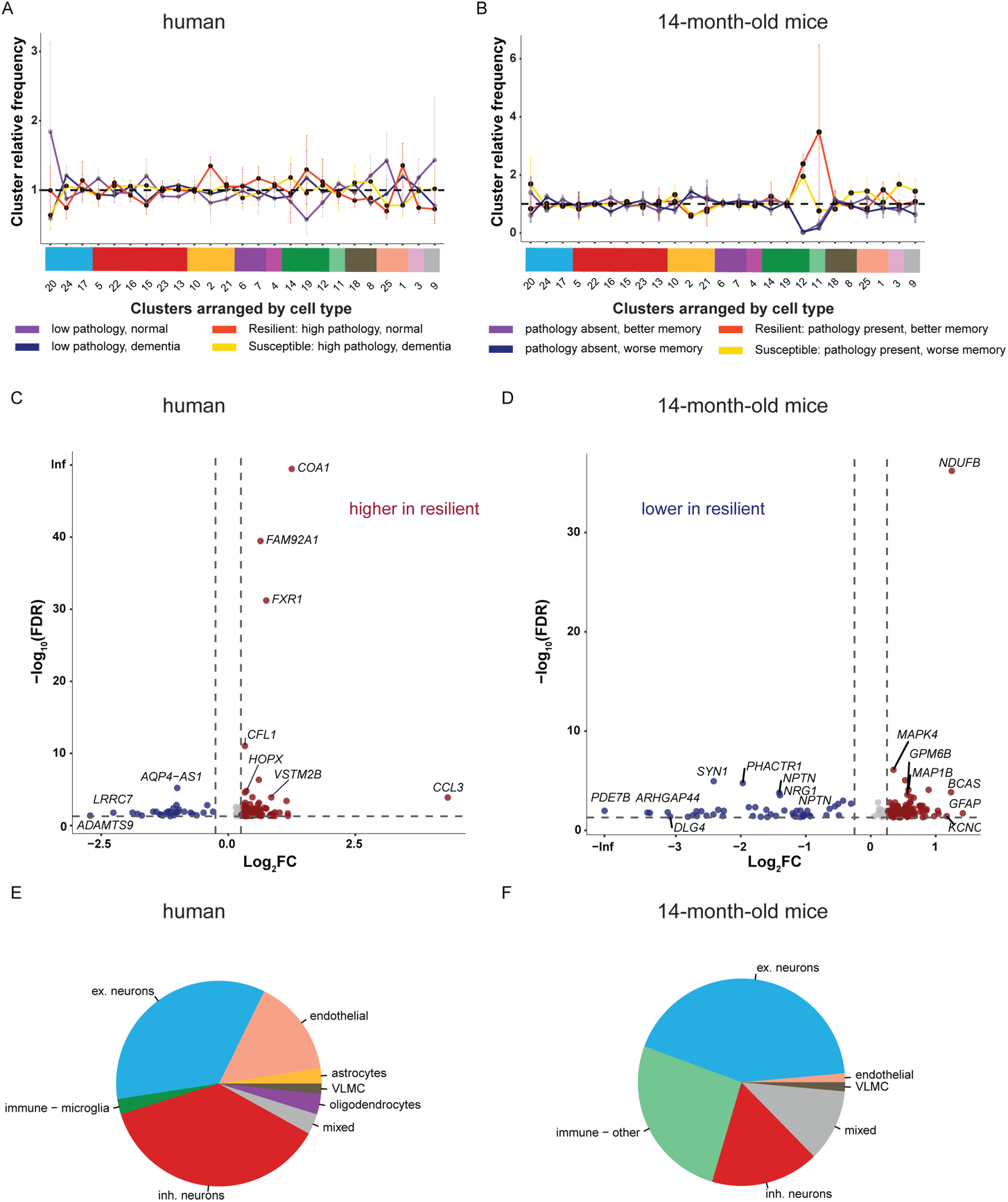
Resilient transcriptional signatures identified in excitatory cluster shared cross-species. **A** and **B**. Composition of all the sample groups in each cluster for human subjects with pathology (**A**) and 14 mo mice with 5XFAD transgene (**B**), with resilient and susceptible groups with more saturated color. No statistical differences were found between the resilient and susceptible groups in each cluster (t-test with Benjamini-Hochberg multiple comparison correction); **C** and **D.** Volcano plots with significantly differentially expressed genes between resilient and susceptible individuals (each dot), for human (**C**) and 14 mo mouse subjects (**D**). Blue color indicates downregulation in resilient individuals (log_2_FC ≤ −0.25) and red color indicates upregulation in resilient individuals (log_2_FC ≥ 0.25); **E** and **F**. Pie charts demonstrating cell type identity of clusters for differentially expressed genes from **C** and **D**. In both in human (**E**) and mouse (**F**), most genes are from neuronal clusters.

Overall, there were 124 human and 142 mouse genes across all clusters that were differentially expressed between resilient and susceptible groups (see Methods, adjusted p-value ≤ 0.05 and log2FC ≥ 0.25 or ≤ −0.25) (Fig. 3C and D), with no overlap between species (Table S3), with most of the differentially expressed genes present in neuronal clusters (Fig. 3E and F). Most downregulated genes in the human resilient group were in the endothelial cluster (C1) and in the mouse in the immune-other (C11) cluster (Figs. 4A and B). Excitatory neuronal cluster 20, annotated as Layer 4/5 intratelencephalic (IT) neurons using MetaNeighbor (Table S2) and confirmed to contain layer-specific genes^15, 16^ (Fig. S2B), contained most of the genes that were significantly differentially expressed and were upregulated in both the mouse and the human subsets (Fig. 4A and B). This neuronal subtype may correspond to a previously reported cell cluster whose signature was associated with residual cognition in bulk human RNA-seq data,^7^ although the cell cluster definition there was based on a healthy donor reference snRNA-seq data set.

**Fig. 4.**
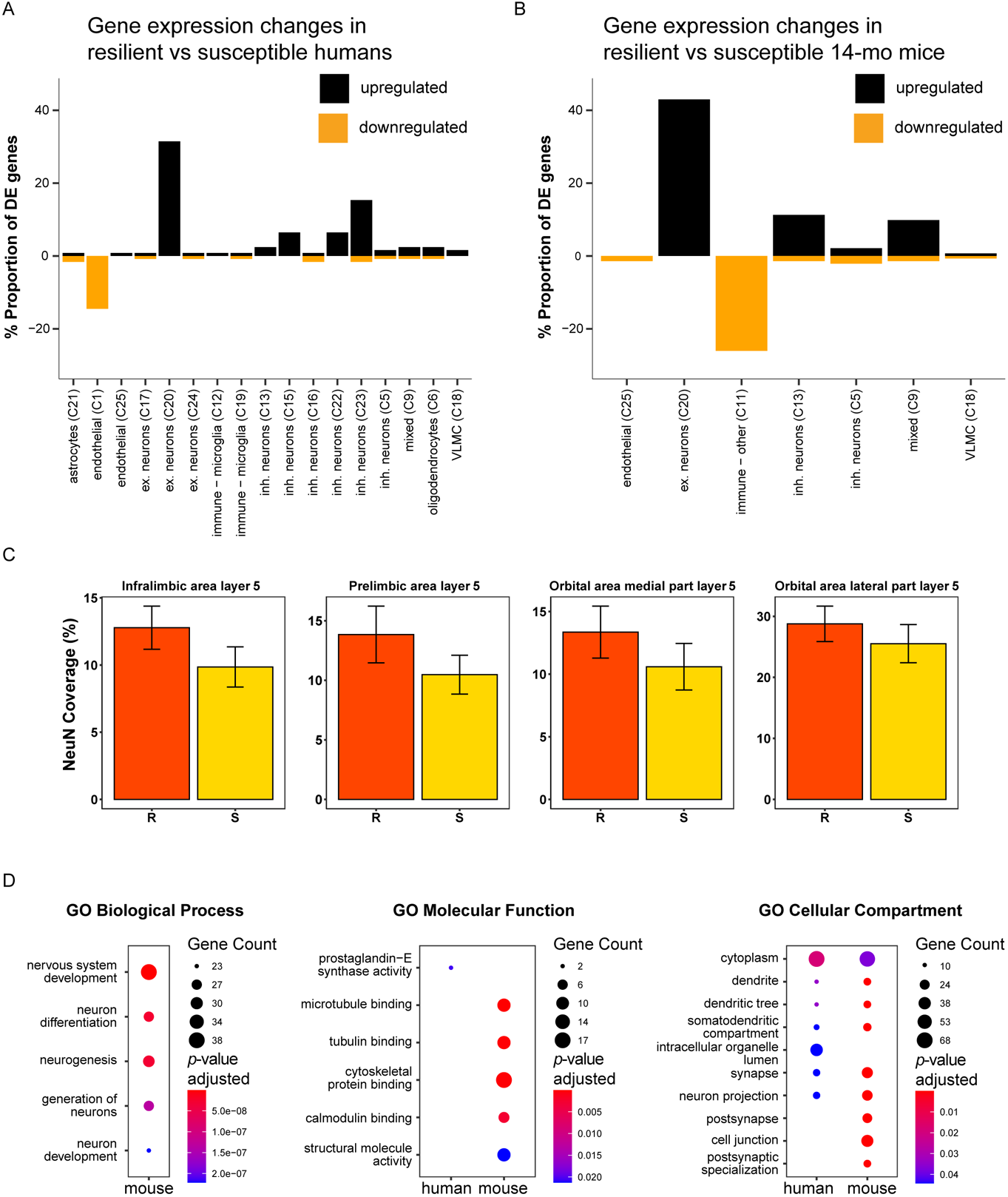
Resilience gene network characterization. **A** and **B.** Proportional significantly differentially expressed (adjusted p-value ≤ 0.05) upregulated and downregulated genes in the human subjects with pathology (**A**) and the 14 mo mice with 5XFAD transgene (**B**) showing most genes were upregulated (log_2_FC ≥ 0.25) and are particularly enriched in cluster 20. Only clusters with significantly differentially expressed genes are shown; **C.** Bar graphs of NeuN coverage in multiple layer 5 regions (orbital area, prelimbic area, and infralimbic area) demonstrate no difference between resilient and susceptible strains. Values with 0% coverage due to absence of area in sections were removed (see also Fig. S2C); **D.** GO enrichment analyses for human and mouse gene lists identified pathways in the mouse data set (5 top pathways included for each species, adjusted p-value threshold of 0.05).

To test the hypothesis that cognitive status was not associated with differences in neuronal composition suggested by comparable relative proportions of nuclei in susceptible and resilient AD strains (Fig. 3A), we quantified the coverage of NeuN positive cells in layers 4 and 5 of the frontal cortex in a separate cohort of AD-BXD mice by immunohistochemistry (IHC). We found that cognitive status was not correlated with regional differences in NeuN load in a population of male and female 14-month-old AD-BXDs. This lack of relationship between neuron coverage within layers 4 and 5 of the frontal cortex and resilient versus susceptible status suggests that gross changes in neuronal cell composition are unlikely to be driving the differences in CFM among the strains assessed (Fig. S2C for correlations on all assessed layer 4 and 5 regions and Figs. 4C for data from selected few layer 5 regions). The same was found when analyzing the female subset of mice; there was no significant association across all frontal cortical regions measured except in the orbital area, medial part, layer 5 NeuN (*p*-value 0.004), but it did not survive FDR correction (*p*-value 0.1) (data not shown). In addition, we confirmed the finding that cluster 20 (layers 4/5) proportions were not different between resilient and susceptible groups in an independent AD-BXD cohort. Deconvolution^17^ of frontal cortex bulk RNA-seq estimated proportions of clusters based on snRNA-seq data (Fig. S2D). Considering each cluster proportion, no statistically significant differences between resilient and susceptible groups of mice were found, including in cluster 20 (Fig. S2D). Therefore, since we observed no change in neuron load by nuclear fraction (Figs. 3B and S2D) and IHC analyses (Fig. 4C), we interpret this as cognitive resilience is conferred, in part, by changes in gene expression levels in layer 4/5 neurons, as there is no observable difference in the degree of neurodegeneration between susceptible and resilient individuals using two orthogonal analyses.

Notably, differentially expressed genes were overrepresented in excitatory neuronal cluster 20. Specifically, our analysis identified 39 human and 61 mouse genes with greater mean and percent nucleus expression in resilient individuals, indicating upregulation may be driven by more neurons expressing these genes. The 61 upregulated mouse genes were not significantly differentially expressed in animals from the corresponding non-transgenic strains at 6 or 14 months or the transgenic animals at 6 months, with one exception that *KCNH7* is downregulated in cluster 20 with log_2_FC −0.51 in 14 months NTG group in resilient group compared to the susceptible one. This suggests that genes are activated in response to the presence of amyloid and increasing age in resilient strains.

When we compared GO enrichment analyses on human and mouse differential gene sets from all clusters, we found that of the two species, only mouse gene GO terms reached significance for biological processes (GO:BP), including neurogenesis, nervous system development, neuron differentiation, generation of neurons, and neuron development (Fig. 4D, far left). For molecular function enrichment, the human gene set list was enriched only for prostaglandin-E synthase activity, while the mouse gene set list was enriched in multiple binding categories, such as microtubule, tubulin, cytoskeletal protein, and calmodulin (Fig. 4D, middle). Genes from both species were enriched for neuron-specific compartments, such as dendrite, synapse, and neuron projection (Fig. 4D, far right). The singular enriched human GO molecular function or biological process pathways is likely due to the fact that there are fewer differentially expressed genes, and poorer gene coverage from the Chromium v2 library for the human compared to the v3 library for the mouse indicated by more reads per nucleus on average in each cluster for mouse than human (mean of average counts per nucleus in each cluster: 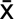 = 0.146 counts, sd = 0.090 counts for mouse; 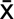 = 0.047 counts, sd = 0.019 counts for human). Additionally, other factors such as longer post-mortem interval during human brain sample collection, as well as the general diversity of human samples due to cumulative lifetime exposure and experiences, may contribute to variability. Since the mouse data set had a greater read depth, we examined the resilience mouse gene list to select resilience transcriptomic candidates from the 14-month-old transgenic cohort in the next sections of this report, focusing on genes in cluster 20. Because of the high alignment of human and mouse transcriptional profiles, we hypothesize that the translational relevance of the mouse resilience findings is high, which we test directly in orthogonal human data sets, below.

### Resilience gene candidates involved in nervous system development and transport pathways

Some genes from mouse cluster 20 that are among the highest ranked top 10 biological processes (in three broad parent categories of nervous system development, transport, and axo-dendritic transport) share pathways (Fig. 5A). For example, 18 genes, e.g. *KALRN*, *GPM6B*, *SPTBN1*, and *PPP3C*, are in the categories nervous system development and transport, while genes *MAP1A*, *MAP2*, and *PAFAH1B1* are in the categories nervous system development and axo-dendritic transport. We also note the different expression profiles in each strain that led to overall resilience profiles in nervous system development (Fig. 5B) and transport genes (Fig. 5C), highlighting the strength of using a genetically diverse mouse panel. Note that gene names were left in the human notation (all letters capitalized) due to the use of human annotations for GO analysis.

**Fig. 5.**
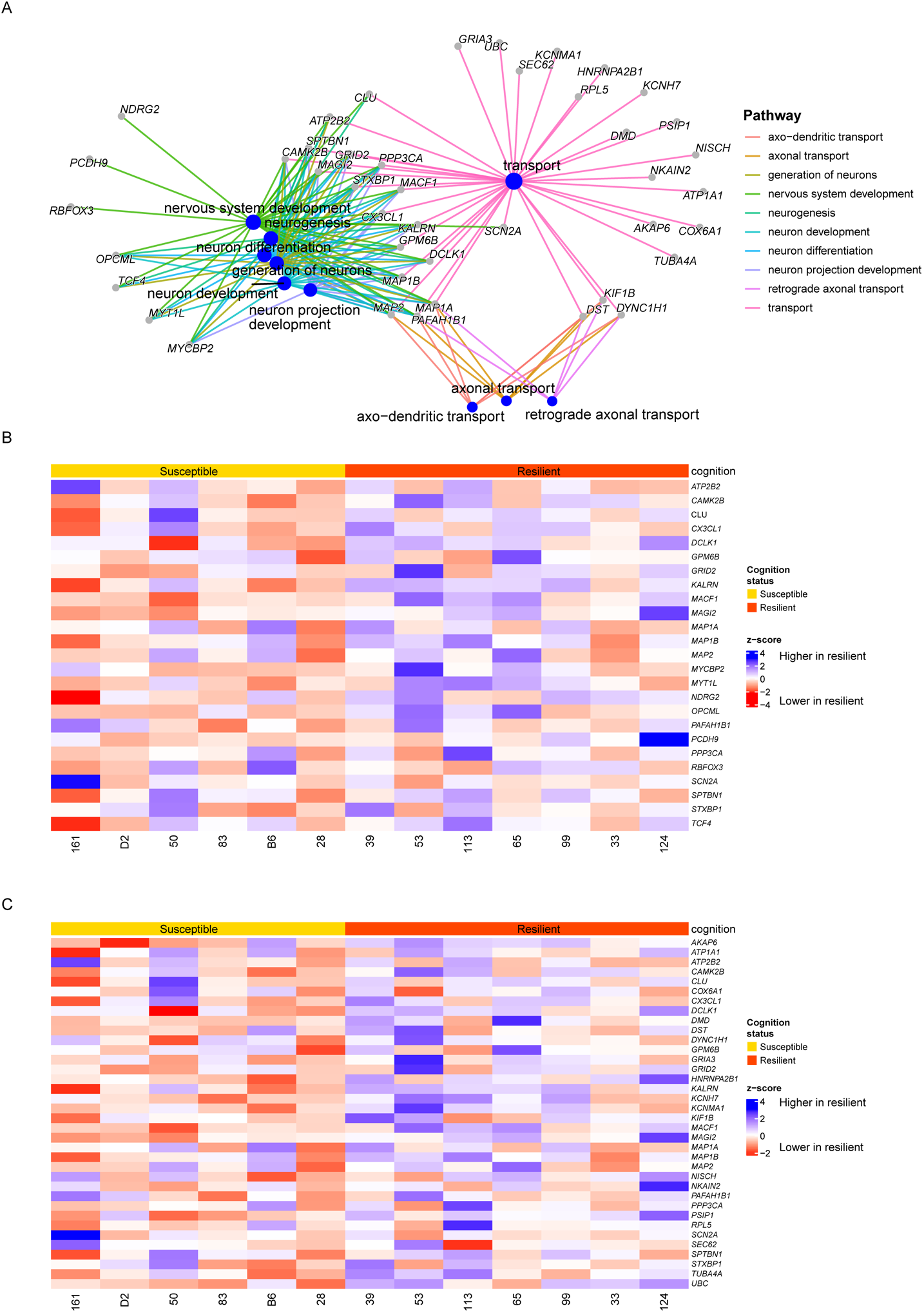
Mouse resilience gene characterization. **A.** Gene concept network demonstrating gene linkages among the top 10 GO:BP terms demonstrating overlap and differentiation of gene function: parent pathways nervous system development (25 genes), transport (36 genes), axo-dendritic transport (6 genes) (see also Figs. S3A, S4A-B). **B.** Heatmap of average normalized gene expression (scaled by gene) for nervous system development pathway, with blue indicating upregulation in resilient mice and red downregulation. **C.** Heatmap of average normalized gene expression (scaled by gene) for transport pathway, with blue indicating upregulation in resilient mice and red downregulation.

Notably, neurogenesis pathway (GO: 0022008) is a top 5 pathway with 21 genes (e.g. *CAMK2B*, *DCLK1*, *MAP1A*, *MAP1B*, *MAP2*, *MACF1, and MYT1L*). Although neurogenesis has been most extensively studied in the hippocampus and the subventricular zone, it has also been observed in multiple other regions in rodent studies.^18^ While the degree to which neurogenesis is present in the adult human hippocampus remains highly controversial,^19–21^ it has been shown to be decreased in neurodegenerative diseases^22^ and associated with cognition.^23^ We observed enrichment of these genes in these resilient cases, although all these genes are also part of the nervous system development process (Figs. 5A and S3A) because the neurogenesis GO term is a child of the nervous system development GO term (GO:0007399). Additional analyses revealed low expression of neurogenesis markers observed in other studies, e.g. *Dcx*, *Mki67*, *Mcm2*, *Lpar1*, *Pax6* and *Sfrp1*^19, 24, 25^ (Fig. S3B). We observed *Dcx* in inhibitory neurons (Fig. S3B), as previously described in the hippocampus.^21^ Thus, it remains unsettled whether the resilience signature in excitatory neuronal cluster 20 is truly indicative of newly born neurons in the cortex. Future work is required to test the prediction that newly born neurons are part of this resilience signature in the cortex.

To further understand known neuronal gene functions associated with resilience in our study, we further classified genes under the broad term “nervous system development” (25 genes). More specific roles for genes were uncovered, including neuron differentiation (*DCLK1*, *GRID2*, *MAP2*, *MYCBP2*, *PAFAH1B1*), axonogenesis (*DCLK1*, *KALRN*, *MACF1*, *MAP1A*, *MAP1B*, *MAP2*, *MYCBP2*, *PAFAH1B1*, *SPTBN1*, *STXBP1*), dendrite morphogenesis (*CAMK2B*, *DCLK1*, *MAP2*, *PPP3CA*), and regulation of synaptic plasticity (*CAMK2B*, *CX3CL1*, *GRID2*, *MAP1A*, *MAP1B*, *STXBP1*) (Fig. S4A). Some of these genes have been previously studied in the context of AD. For example, proteins MAP1A, MAP1B, and MAP2 are microtubule-associated proteins with function in microtubule assembly during neurite growth and morphogenesis, known to interact with MAPT (microtubule-associated protein tau), and have been shown to be involved in AD pathogenesis.^26, 27^ *DCLK1* codes for another microtubule-associated protein, Doublecortin Like Kinase 1, that regulates neurogenesis.^28^ *PPP3CA* (Protein Phosphatase 3 Catalytic Subunit Alpha), involved in presynaptic and postsynaptic phosphorylation, had different pattern of isoform expression in AD vs controls.^29^ *CX3CL1* codes for a chemokine Fractalkine, a ligand for CX3CR1, enabling interaction between neurons and microglia^30^ that was found to have a role in neurogenesis.^31^ It was also reported that the hippocampal levels of *CX3CL1* are lower in late AD stages.^32^ Our findings add to the growing body of evidence that resilience is associated with improved neuronal function.^9^ Yet, how transcriptomic upregulation of these genes translates to improved cognition remains to be discovered.

Many genes in the broad category “transport” (36 genes) that we identified to be upregulated in resilient mice have known functions in the central nervous system. More narrowly defined enriched pathways include vesicle and organelle transport along microtubules and regulation of membrane potential (Fig. S4B). Genes categorized as involved in microtubule transport include *DST* (Dystonin), which codes for the BPAG1 protein, a linking cytoskeletal protein^33^ and member of the plakins family of gigantic crosslinking proteins that include another resilience upregulated gene, *MACF1*^34^; *DYNC1H1* (Dynein Cytoplasmic 1 Heavy Chain 1), a heavy chain of a motor complex whose mutations are associated with various neurological diseases by reducing the range of complex movement along microtubules^35^; and *KIF1B* (Kinesin Family Member 1B) that codes for a motor protein whose mutation causes Charcot-Marie-Tooth disease.^36^ Regulation of membrane potential include *ATP1A1 (*ATPase Na+/K+ Transporting Subunit Alpha 1), coding for a protein with a role in establishing sodium and potassium gradients; *DMD* (Dystrophin), mutations of which cause Duchenne muscular dystrophy; and G*RID2* (Glutamate Ionotropic Receptor Delta Type Subunit 2). The fact that these genes were upregulated in the resilient mouse group may mean that microtubule transportation processes are retained. In fact, genes whose function involves axonal guidance (*UNC5C*) and acting binding (*ENC1*) were previously identified as resilience factors.^37^ As well, neuronal transport deficits have been identified in AD,^38, 39^ and while microtubule stabilization reduced pathology and improved cognition in AD mouse models,^40–42^ challenges remain to translate these findings to the clinic.^43, 44^ The remaining parent pathway – “axo-dendritic transport” – has overlapping genes with the previously discussed pathways and provides further biological processes likely involved in resilience. Overall, biological processes pathways enriched in these genes point to an improved ability to move cargo along microtubules in neurons. Alternatively, neurons in the resilient mice may be undergoing processes to remodel the cytoskeleton to retain functionality under stress.^45^ The upregulation of these genes involved in transport in resilient individuals points to overall improvement of neural function.

### Downregulated genes regulate neuronal processes

Additionally, the clusters containing most downregulated genes in the human (endothelial C1) and in the mouse (immune - other C11) were identified. For human downregulated genes, 18 of the 30 total downregulated genes were in endothelial cluster C1, including *DPP10*, *ENO2*, and *MAGI2*. GO Biological Processes analysis revealed multiple pathways related to neuronal function, such as regulation of neuron projection development (*LRRC7*, *MAGI2*, *MAP3K13*, *NEGR1*) (Fig. S5A). Endothelial cells have been shown to influence neurogenesis and axonal growth through release of secreted factors,^46, 47^ yet, how these downregulated genes are involved in resilience requires further investigation. Interestingly, gene *AQP4-AS1*, a long non-coding RNA, was downregulated in three clusters: inh. neurons C16 and C23, and ex. neurons C24 (Table S3). For the mouse, 37 of 47 total downregulated genes were in the immune – other cluster 11 with lymphocyte marker genes *CCL5* and *NKG7*.^48, 49^ Genes in this cluster are also involved in neuronal function, such as synaptic signaling and plasticity, neuron projections and differentiation (Fig. S5B). Lymphocytes are known to be involved in AD; for instance, T cells infiltrate the CNS in AD with a potential role in disease pathology^50^ and directly interact with neurons and promote neuroinflammation.^51^ Due to the small size of the clusters C1 and C11, it is difficult to interpret these findings in the context of resilience. Further studies with enrichment of these cell types would be necessary to determine their role in resilience.

### Resilience candidates are upregulated in excitatory neurons and correlate with cognition in independent human reference data sets

We found corroborating evidence of these upregulated mouse genes among several human data sets. Remarkably, 25 genes were predicted to have differential expression in resilient individuals in various tissues, including the brain, using PrediXcan analysis in GWAS of resilience (Table S4) and expression of 38 genes across three tissues, including the DLPFC, in bulk transcriptomic data were found to be positively correlated with cognition (Table S5).

When considering the products of the 61 genes assessed, 23 proteins with significant positive correlations between protein abundance and MMSE (mini-mental state exam) (p < 0.05; higher abundance correlated with better cognition) (Table S6) were identified in the ROSMAP and Banner (Banner Sun Health Research Institute) populations.^52^ Johnson et al reported that modules “post-synaptic density” and “protein transport” were enriched in proteins correlated with resilience that were previously identified in another study.^52^ In an analysis that directly compared protein abundance in AD vs AsymAD (individuals with AD pathology and without significant cognitive impairment) (Supplementary Table 2 in Johnson et al), 1,540 proteins had higher abundance in AsymAD (log_2_FC > 0 with adj. p-value < 0.05). Our results enable narrowing down of resilience factors and nomination of cell-type specific targets. In fact, we matched 19 of our gene targets with the protein list, including ATP1A1, ATP2B2, DCLK1, GRIA3, KCNMA1, KIF1B, MAGI2, MAP2, SPTBN, and STXBP1. Overall, these gene expression and protein data cross-check strengthen confidence that the resilience factors that were identified in the mouse excitatory neuronal cluster 20 from the cross-species integrated data set are translationally relevant.

### Nominated resilience candidates include druggable targets

An important factor to consider in nominating targets is whether the gene is druggable. Our study has nominated genes that are targetable based on Agora’s druggability criteria (https://www.synapse.org/#!Synapse:syn13363443). After obtaining these data for each gene, we filtered the list by druggability, safety, and accessibility of target. Those genes that are targetable by small molecules, homology, or structure were retained. Of the genes that are targeted by protein structure, only those with two or fewer “red flags” such as it being an essential gene, a cancer driver, having high off target gene expression, or a “black box warning” (label put on by the U.S. FDA to warn about serious safety risk) on drugs currently used in the clinic, were retained. Finally, we arrived at a list of 17 targetable genes (Table S7), six of which are targetable by small molecules and have a favorable safety profile: *ATP1A1*, *GRIA3*, *KCNH7*, *NISCH*, *SCN2A*, and *TUBA4A*.

In addition, we reviewed which of the 61 genes had been previously nominated as potential targets for AD. Among the upregulated resilience genes from excitatory neuronal cluster 20, 11 mouse genes have been nominated on Agora by research teams from the NIA-funded AMP-AD consortium (https://agora.adknowledgeportal.org/genes) (Table S8). None of these targets have been validated and one target – *SCN2A* – is subject of ongoing validation studies. Our data provide new evidence to prioritize these genes for candidate validation: *ATP2B2*, *CLU*, *CX3CL1*, *DCLK1*, *GFAP*, *GPM6B*, *KALRN*, *MAP1B*, *MAP2*, *and PPP3CA*.

### Drug repositioning candidates identified to promote resilience motifs

Using a complimentary strategy to nominate resilience drug candidates to test, we implemented a tiered computational approach to identify compounds that are most likely to boost the resilience signature (or “motifs”) by increasing expression of multiple target genes. Accordingly, we matched the 61-gene resilience signature with patterns of transcriptional changes upon drug administration to different cell lines (see Methods). Through this process, we identified 93 drugs with known biological function for potential repurposing to increase cognitive resilience in AD (Table S9). Moreover, we highlight nine drug candidates that stood out beyond the rest based on highest score (Table S9, global score of 0.9 or above), and we nominate them for priority testing: velnacrine, miglitol, sirolimus, BRD-K63175663, W-13, methotrexate, Mebendazole, podofilox, and BMS-536924. Interestingly, the top scorer is velnacrine, a synthetic AChE inhibitor that was considered for treatment of AD but discontinued.^53^ Other AChE interactors outside of top candidates were an antimalarial mefloquine (score 0.24) and a withdrawn drug for ulcers – ranitidine (score 0.15). Notably, rapamycin (synonym sirolimus), an MTOR inhibitor with efficacy in animal AD models^54^ and currently in clinical trial phase 2 (NCT04629495), is a top candidate with score of 0.98. As well, rapamycin acts on FGF2, a previously nominated modulator of short term memory by our group.^12^ Other notables in the top nine are tubulin and microtubule inhibitors mebendazole and podofilox. Interestingly, drug bepridil, an ATP1A1 inhibitor, is also on the list with score of 0.11 (Table S9). To gain further biological insight into the 93 drugs considered for repurposing, GO Enrichment analysis on the known target genes of those drugs (also referred to as drug set enrichment analysis) was performed. We identified shared biological processes pathways related to neurotransmitter uptake, dopamine and catecholamine in particular (Table S10), with nitric-oxide synthase and catecholamine binding molecular functions enrichment (Table S11), and cellular compartment enrichment at the synapse with terms of dopaminergic and GABA-ergic synapse, potassium and calcium channel complexes, as well as spindle microtubule among the top terms (Table S12).

## Discussion

Our findings demonstrate that analysis of integrated mouse and human transcriptomic data uncovers novel and conserved cell type-specific signatures in layer 4/5 cortical neurons associated with resilience to AD cognitive decline. The integrated approach ensures same treatment of data and allows for straightforward mouse to human comparisons. Using this approach, we identified resilience genes that were confirmed in other human data sets. Specifically, we demonstrated that the majority of cell types resolved from mouse and human snRNA-seq data sets exhibited high alignment, which allowed us to identify translationally-relevant resilience signatures conserved across the two species. We determined that the upregulation of gene expression in layer 4/5 excitatory neuron cluster reflects a robust signature of resilience, while cluster composition remains stable. This is related to, but distinct from, a recent report implicating a RORB+ neuronal cluster signature associated with residual cognition in bulk RNA-seq data from the ROSMAP cohort. In particular, we refined our investigation of this layer 4/5 neuronal cluster by cross-referencing human and mouse data, and also investigating specific gene expression signatures that are altered within this cluster. Through this process, we nominated targeting of specific genes such as *ATP1A1*, *GRIA3, KCNMA1,* and *STXBP1* in pathways such as regulation of membrane potential and axonal transport in excitatory neurons to promote resilience.

While animal models are necessary for development of AD therapeutics, more translationally-relevant mouse models are needed due to the lack of clinical trial success.^55, 56^ Large-scale efforts to characterize human cohorts and mouse models allows for better understanding of the molecular underpinnings of AD.^57^ In particular, several studies reported both shared and divergent signatures between mouse and human transcriptomic AD signatures in microglia and other cell types.^58–61^ However, many studies share limitations of comparing gene signatures obtained from different computational pipelines, comparing different technology (e.g., snRNA-seq vs scRNA-seq), comparing different brain regions, and only using one or a limited number of genetic backgrounds for mouse models. In our study, we aimed to eliminate several of these technical differences that may confound biological interpretation of results. Therefore, we performed a unified analysis using the same computational pipeline, sequencing technology, and included genetic diversity in profiling a similar region from the mouse and human brain tissue (Fig. 1A). While not all technical differences were eliminated, we demonstrate that this analysis pipeline is a powerful tool to synthesize transcriptomic data into an interpretable cross-species data set (Fig. 1B-E).

We discovered that while cross-species cluster composition remained unchanged with cognition status (Fig. 3A and B), both human and mouse resilient individuals had an upregulation of genes in one excitatory neuronal cluster (Fig. 4A and B) that was classified as layer 4/5 IT neurons (Table S2). IT neurons are a diverse class of neurons (in terms of connections, activity, and morphology) in layers 2-6 that project to telencephalon and contralaterally.^16, 62^ IT neurons in layer 4 process external input, while IT neurons in other layers receive input from L4 and from external sources.^16^ Some signature genes that have been identified in 4/5 IT neurons include *Rorb* and *Satb2.*^16, 62^ Interestingly, excitatory neurons expressing RORB were found to be vulnerable in the entorhinal cortex region in human AD,^63^ and also associated with residual cognition in bulk RNA-seq data analysis.^7^ Thus, converging lines of evidence suggest that targeting excitatory IT neurons in these layers may be a therapeutic strategy to engage resilience mechanisms. We note that the resilience-associated genes within this cell group are not the same in human versus mouse; this could be due to differences in species, genetics (familial AD mutations in mouse versus late-onset AD in human individuals), pathology (amyloid in mouse versus amyloid and tau in human), or variation in our relatively small sample of humans. However, the fact that the same neuronal group is implicated in both species *despite all of these differences* is highly significant.

In the approach to targeting excitatory neurons to promote resilience, our data indicate that it is the gene expression that matters, not the survival of neurons. First, we observed that there was no difference in cluster composition (at our optimal cluster resolution) between resilient and susceptible groups (Fig. 3A and B). Second, we corroborated this finding by IHC analysis of NeuN load in layers 4 and 5 of the frontal cortex in the mouse (Figs. 4C and S2C) as well as in a different cohort of AD-BXD mice (Fig. S2D). While the 5XFAD mutation itself resulted in a decrease of the number of neurons in layer 5 in middle age mice on C57Bl/6 x SJL and C57Bl/6J backgrounds,^64, 65^ the difference in neuronal proportions between resilient and susceptible individuals is less clear. In a ROS cohort, same density of neurons was found in AD-resilient (AD pathology and cognitively normal) and AD (AD pathology and dementia) individuals in midfrontal gyrus cortex (frontal lobe).^6^ While some coverage analyses show no difference in neuronal load, they do not take into account brain region shrinkage that is typical in AD. In a study where neurons were counted, fewer neurons were observed in AD patients compared with people with AD pathology but no cognitive symptoms in superior temporal sulcus area (temporal lobe).^66^ In the hippocampus, some regions maintained neuronal numbers while others showed decline between pre-clinical AD (cognitively normal people with AD pathology) and AD in Baltimore Longitudinal Study of Aging and the Johns Hopkins Alzheimer’s Disease Research Center cohorts.^67^

A closer examination of differentially expressed genes within this IT neuronal cluster revealed potential biological processes of resilience. The broad parent biological processes of the 61 mouse genes were nervous system development, transport, axo-dendritic transport (Fig. 5A). While there were 21 genes in the broad category of neurogenesis, we observed no neurogenesis signature in cluster 20 (Fig. S3B). The genes that were labeled as part of the nervous system development were further classified by more specialized categories such as involvement in neuron migration, axonogenesis, regulation of synaptic plasticity and trans-synaptic signaling, and dendrite morphogenesis (Fig. S4A). Activating these pathways may improve resilience to AD cognitive decline. Additionally, we identified genes in pathways transport and axo-dendritic transport to be upregulated (Figs. S3A and S4B) and hypothesize that resilient individuals are able to retain essential neuronal function, as this signature appears late in transgenic animals. Our hypothesis is bolstered by other studies showing that overexpression of a microtubule motor protein *Kif11* improved cognition in 5XFAD mouse model while not changing Aβ load.^68^

To provide additional evidence of translatability of the resilience genes identified in the mouse, we cross-referenced our findings with a different human cohort of resilient individuals and/or data modality. Remarkably, we found corroborating evidence among many of the 61 mouse genes in human cohort transcriptional data sets, including predicted differential expression in resilience in multiple tissues (Table S4) and positive correlation with cognition (Table S5). As well, we found abundance of over a third of proteins that are products of the resilience genes to be positively correlated with cognition (Table S6).

Finally, in nominating resilience targets we considered the druggablity of these genes. From mining Agora druggability data set, we tabulated targets with the best druggability metrics. From the original list of mouse genes, 11 mouse genes have been nominated on Agora (Table S7), but only *SCN2A* and *MAP1B* had favorable druggability profiles, which is why studies such as this are required to nominate more targets. We found 16 additional druggable genes among those that hadn’t been previously nominated, including seven that were validated across orthogonal transcriptomic and proteomic human datasets: *ATP1A1*, *DCLK1*, *GRIA3*, *KCNMA1*, *MAP2*, *STRBP*, and *STXBP1*.

To highlight four of these seven targets, we will discuss *ATP1A1*, *GRIA3, KCNMA1, and STXBP1* in some detail. *ATP1A1* (ATPase Na+/K+ Transporting Subunit Alpha 1) codes for one of the four that are catalytic α subunits of Na^+^/K^+^ ATPase, which is involved in resting membrane potential; its mRNA is expressed in neurons throughout the mouse brain.^69^ *GRIA3* (Glutamate Ionotropic Receptor AMPA Type Subunit 3) codes for protein Glutamate Receptor 3, a subunit of the AMPA receptor, with function in synaptic transmission. Mutation in this gene has been associated with cognitive impairment,^70^ and levels of GluR3 have been found to vary during AD progression.^71, 72^ *KCNMA1* (Potassium Calcium-Activated Channel Subfamily M Alpha 1) that codes for a subunit for a calcium activated big potassium channel with a role in neurotransmitter release. *KCNMA1* SNPs were found to be associated with age at onset and duration of AD^73^ as well as increasing odds ratio of sporadic AD.^74^ *STXBP1* (Syntaxin Binding Protein) codes for protein MUNC18-1 that interacts with the SNARE complex involved in neurotransmitter release that has recently been found to interact with tau.^75^ Another SNARE-related protein, CPLX1, had been previously shown to be associated with cognitive resilience.^4^ Interestingly, mRNA levels of *KCNM1*^74^ and *ATP1A1*^76^ have been reported to be elevated in AD. In our mouse snRNA-seq data, these four genes are expressed in at least 10% of nuclei in both groups of mice, with proportionally more resilient nuclei expressing them, while in nuclei that do express the gene, the average expression between resilient and susceptible groups is similar. Therefore, we postulate that upregulating gene expression of these genes in excitatory layer 4/5 neurons may result in improved cognitive resilience.

Small molecules that have been approved for some of these genes, listed on genecards.org, include both agonists and antagonists. Focusing on agonists to potentially upregulate cognitive resilience, we identified several potential candidates to test in *in vitro* and animal models. For instance, approved small molecule agonists for *ATP1A1* include magnesium gluconate used for hypomagnesemia (https://go.drugbank.com/drugs/DB13749), for *GRIA3* include cyclothiazide, a diuretic (https://go.drugbank.com/drugs/ DB00606) and venlafaxine, a serotonin and norepinephrine reuptake inhibitor (https://go.drugbank.com/drugs/DB00285), and for *KCNMA1* another diuretic bendroflumethiazide (https://go.drugbank.com/drugs/DB00436). On the other hand, STXBP1 does not have an approved drug agonist. For drug targets lacking agonists, novel small molecules or gene therapies are needed for validation and pre-clinical studies. Notably, venlafaxine was also on the list of drugs for repurposing that upregulate resilience motifs with a score of 0.24 (Table S9), demonstrating that multiple drug nomination routes converge on same drugs. Our drug repurposing approach leverages the known properties of compounds that can be applied in the field of resilience. We identified other candidates that increase the resilience motifs that function in the CSN and at the synapse (Tables S10-12), with top candidates including rapamycin and velnacrine (Table S9). Interestingly, in our 5XFAD amyloid model of AD, we identified microtubule-associated resilience gene candidates and some of our top candidates for drug repurposing are mebendazole and podofilox, tubulin inhibitors. It has been recently published that mouse genetic background modulates mutant tau pathogenicity.^77^ Whether targeting these genes specifically in excitatory neurons in layers 4/5 enhances resilience is a topic of future investigations.

It is worth noting that in ROS/MAP and other cohorts, researchers have previously identified factors such as lifestyle and complex traits,^5, 78^ pathology burden,^79^ cellular and structural markers,^6, 66, 80^ genes,^37, 81^ and proteins^6^ associated or correlated with resilience (recently reviewed in ^9^). However, proving causality and translating findings to therapeutics remains a challenge. Using the translationally-relevant AD-BXDs, in future studies we aim to study causality, understand the mechanisms of resilience that we nominated in the genetically diverse mouse models, as well as to perform preclinical studies to test top candidates.

In conclusion, our cross-species integrative transcriptomic analyses of individuals resilient to AD cognitive decline resulted in nomination of several resilience targets. The AD-BXD mouse panel allows us to decipher causality and perform mechanistic and pre-clinical animal studies in forthcoming projects.

## Acknowledgements

We gratefully acknowledge the contribution of Mike Samuels, Shannon Bessonett, and the Single Cell Biology service, the Genome Technologies service, and Advanced Cyberinfrastructure high performance computing resources at The Jackson Laboratory for expert assistance with the work described in this publication. We acknowledge Stephan Fischer’s role in MetaNeighbor studies. The results published here are in part based on data obtained from Agora, a platform initially developed by the NIA-funded AMP-AD consortium that shares evidence in support of AD target discovery (https://agora.adknowledgeportal.org/genes, Site Version 2.2.0-3a1674d, Data Version syn13363290-v33). We are thankful for the funding for this project: R01 AG057914 (C.C.K.); R01 AG054180 (C.C.K); R01 AG075818 (C.C.K.), RF1 AG059778 (C.C.K), Alzheimer’s Association Zenith Fellows Award AARF-ZEN-21-846037 (C.C.K. and M.A.T.); R01AG059716 (T.J.H.), R01AG061518 (T.J.H.), R01AG074012 (T.J.H.), R01 AG066831 (V.M.); RF1 AG057473 (V.M.); U54 AG076040 (V.M.); Thompson Family Foundation TAME-AD Award (V.M.); U01AG061357 (N.T.S.); Swiss National Science Foundation fellowship P2EZP3_191873 (S.M.); R01LM012736 (J.G.); R01MH113005 (J.G.); RF1AG058942 (M.A.); RF1AG059093 (M.A.); U01AG061359 (M.A.); U19AG063744 (M.A.); and R01AG069901 (M.A. and O.B.). ROSMAP is supported by P30AG10161, P30AG72975, R01AG15819, R01AG17917. U01AG46152, U01AG61356. ROSMAP resources can be requested at https://www.radc.rush.edu.

## Author contributions

Conceptualization, C.C.K, V.M., M.A.T.; Methodology, M.A.T., N.H., Y.D., J.A.L.W., J.-G.Z., M.A., T.J.H., V.M.P., V.M., C.C.K.; Software, N.H., J.-G.Z., V.M.P., Y.D., J.A.L.W, S.M., J.G.; Validation, B.G., Y.W., L.D., E.B.D., N.T.S., T.J.S.; Formal Analysis, M.A.T., B.G., Y.W., L.D., T.J.H., O.B., M.A; Investigation, M.A.T., Y.D., A.R.O., S.M.N., O.B., M.A., J.-G.Z., J.A.L.W., Y.W., L.D.; Data Curation, A.R.D., Y.D., J.-G.Z.; Resources, P.R., S.M., J.G., E.B.D., N.T.S.; Writing – Original Draft, M.A.T., C.C.K, V.M., B.G.; Writing – Review & Editing, P.R., J.-G.Z., S.M., N.H., A.R.O., B.G., M.A., K.M.S.O., C.C.K., V.M., M.A.T.; Funding Acquisition, C.C.K., V.M., T.J.H., N.T.S., J.G., M.A.; Supervision, V.M., C.C.K., P.R., T.H.J., J.G., V.M.P., M.A.

## Methods

### Mouse subjects

The AD-BXD panel of mice were generated as previously described.^10^ Briefly, female 5XFAD mice on C57BL/6J background harboring five human mutations that cause familial AD (Stock No: 34848-JAX; B6.Cg-Tg(APPSwFlLon,PSEN1*M146L*L286V)6799Vas/Mmjax) were mated with males from the BXD panel.^82–84^ The resulting F1 mice were group-housed (2-5 mice/cage) in a facility with a 12-hour light and dark cycle and had free access to food and water. Only female mice were used in this study. Mouse studies were carried out at the Jackson Laboratory and the University of Tennessee Health Science Center and were approved by the Institutional Animal Care and Use Committee (IACUC) at each location. All animal studies were conducted in compliance with the National Institutes of Health Guidelines for the Care and Use of Laboratory Animals.

### Contextual fear conditioning

Mice were trained on contextual fear conditioning (CFC) paradigm as reported previously ^10, 85, 86^. After three days of habituation to transport and to the testing room, the mice were placed in a testing chamber. After a brief baseline period (150-180 s), four mild foot shocks (1 second, 0.9 mA) separated by 115±20 seconds were applied. Contextual fear memory (CFM) was measured 24 hours later by placing the mouse in the same testing chamber and measuring percent freezing during a 10-minute time period. Female AD-BXDs and their non-transgenic littermates were tested on CFC at 6 and 14 months of age.

### Selection of strains for transcriptomic analyses

We analyzed a group of 14 strains, including two founder strains (B6*B6 and B6*D2 F1s) that differed in CFM at 14 months. Strains were stratified into resilient and susceptible based on population mean. AD-BXD strains were 28, 33, 39, 50, 53, 65, 66, 83, 99, 113, 124, 161, and the B6 and B6*D2 founder strains, along with the non-transgenic counterparts to all 14 strains. To select samples that were most representative in terms of CFM for snRNA-seq analyses, samples from mice nearest the strain average of CFM value (where available) were selected for each strain at 6 (mature) and 14 (middle-aged) months of age for both 5XFAD and their non-littermate controls, resulting in 56 total samples. Quality control was performed on snRNA-seq data as follows: samples that were below 2.5 standard deviation of median genes per nucleus, total genes detected, and median UMI counts were replaced with samples from a mouse with the same demographics (strain, age, 5XFAD mutation status). This resulted in three samples needing to be replaced. Mean ages with standard deviation for each group were as follows: mature adult mice: 5.96 ± 0.44 months for transgenic animals; 6.00 ± 0.49 months for non-transgenic animals; middle-aged mice: 13.98 ± 0.28 for transgenic animals; and 13.92 ± 0.41 months for non-transgenic animals (n = 14 for each group, and n = 7 for cognitively impaired and unimpaired within each group) (Fig. 1A).

### Single nucleus RNA-sequencing for mouse subjects

Mice were anesthetized with isoflurane and decapitated after CFM testing. The brains were removed and dissected after olfactory bulbs removal. Nuclei were isolated from snap-frozen frontal cortex (anterior to the anterior forceps of the corpus callosum) samples from one hemisphere using Nuclei Isolation Kit: Nuclei EZ Prep (Sigma-Aldrich Cat. No. NUC-101). Briefly, 50 µl of EZ Lysis Buffer and RNase inhibitor (1000 units/mL; Protector RNase Inhibitor, MilliporeSigma Cat. No. 03335399001, 2000 U, 40 U/µL) were added to 1.5 mL DNA LoBind tubes (Eppendorf Cat. No. 022431021) containing brain tissue. Tissue was ground with Bel-Art Disposable Pestle (Sigma Aldrich Cat. No. BAF199230000). The tissue was washed off from the pestle with 25 µl of Lysis Buffer, and the tube was placed on ice for 5 minutes. The sample was centrifuged at 500xg for 5 minutes at 4 °C. Supernatant was removed and the pellet was resuspended in 50 µL of Lysis Buffer using a wide bore tip. The sample was placed on ice for 5 minutes, after which 50 µl of PBS containing 0.04% BSA (from LAMPIRE Biological Laboratories Cell Culture Grade 35% BSA Liquid (Fisher Scientific Cat. No. 50-414-159) and RNase inhibitor (40 units/mL) were added. The sample was centrifuged for 5 minutes at 4 °C at 500xg. Thereafter, the sample was resuspended in 100 µl of PBS containing 0.04 % BSA and RNase inhibitor and pushed through a pre-wet 40 µm filter (PluriStrainer Mini 40 µm Cell Strainer (Pluri Select Cat. No. 43-10040-60)). The sample was centrifuged for 5 minutes at 4 °C at 500xg, and resuspended in 100 µl of PBS containing 0.04% BSA and RNase inhibitor, then pushed through a pre-wet 5 µm filter (PluriStrainer Mini 5 µm Cell Strainer (Pluri Select Cat. No. 43-10005-60)). After a final centrifugation step at 500xg for 5 minutes at 4 °C, the sample was resuspended in 1000 µL of PBS containing 0.04% BSA and RNase inhibitor and immediately processed as follows.

Nuclei quality was assessed via brightfield imaging and counted via Trypan Blue and a Countess II automated cell counter (ThermoFisher), and up to 12,000 nuclei were loaded onto one lane of a 10X Chromium Controller. Single nuclei capture, barcoding and library preparation were performed using the 10X Chromium platform^87^ version 3 chemistry and according to the manufacturer’s protocol (#CG00052). cDNA and libraries were checked for quality on Agilent 4200 Tapestation, quantified by KAPA qPCR, and pooled at 33.33% of an Illumina NovaSeq 6000 S2 flow cell lane, targeting 6,000 barcoded nuclei with an average sequencing depth of 100,000 reads per nuclei.

Illumina base call files for all libraries were demultiplexed and converted to FASTQ files using Illumina bcl2fastq 2.20.0.422. A filtered digital gene expression matrix was generated for each gene expression library against the 10X Genomics mm10 reference build (version 3.0.0, GRCm38.93 including introns for pre-mRNA mapping) using 10X Genomics CellRanger count version 3.1.0 for all samples.

### Confirmation of sample identity

After sequencing data for 56 samples was obtained, as part of quality assessment analysis, we checked sample identity to confirm correct sex, transgene status, and strain. Using a strain matching tool RNA-strain-match,^88^ we identified two samples that were prepared and run in sequence had flipped strains, which we attribute to a sample swap during sample prep. We re-assigned correct IDs for these two samples. As well, one other sample transgene status was not concordant with initial genotyping results. Genotyping for that mouse tissue was repeated and it matched sequencing results, leading to reassignment of transgenic status of that mouse. Subsequently, while all samples were used for integration and cell-type assignment analyses, due to the lack of full coverage of genotype at each age point, that strain (66) was removed from downstream analyses of resilience signatures.

### Single nucleus RNA-seq data for human subjects

As described in the original report,^13^ frozen dorsolateral prefrontal cortex (DLPFC) brain tissue was profiled for snRNA-seq using 10X Chromium version 2 chemistry for 24 ROSMAP individuals, with six donors per group (three male and three female) from four categories: cognitively normal with low AD pathology, cognitively normal with high AD pathology (resilient), cognitively impaired with low AD pathology, cognitively impaired with high AD pathology (susceptible) (Fig. 1A). General QC measures are described in the original study, and all data is available on the AD Knowledge Portal hosted on Synapse.

### Mouse data set preparation

The mouse data set was combined from individual files using Seurat package^89, 90^ in R^91^, with features retained when expressed in at least 10 nuclei and nuclei kept with at least 200 features. Then, nuclei with mitochondrial, ribosomal, and pseudo-genes over 5 % and RNA count below 500 or above 20,000 were excluded. Thereafter, all mitochondrial genes were removed. The data set was batch corrected on sequencing date using harmony package^92^ with 30 dimensions and resolution of 0.5. Doublet check was performed with 5 % rate using DoubletFinder R package.^93^ According to pre-set thresholds, three clusters with doublet rates of over 40% and one cluster with under 100 nuclei were removed from further analysis, resulting in 36 clusters for 26,006 genes by 186,900 nuclei.

### Human data set preparation

The raw human gene counts and the metadata^13^ were re-processed to ensure same treatment as the mouse data set. The human data set was taken through an identical pipeline as the mouse data set, with two modifications. One, an additional first step of removing duplicated genes after summing counts was done. Two, no batch correction was performed. According to the pre-set thresholds, one cluster with both high doublet rate and fewer than 100 nuclei was removed, resulting in 22 clusters for 26,805 genes by 168,282 nuclei.

### Integration of Datasets

An average of 7,012 and 3,338 nuclei per sample for human and mouse samples, respectively was obtained after filtering and quality control steps, totaling 168,282 human and 186,900 mouse nuclei.

#### Gene name translation

Mouse gene names were converted into homologous human gene names using curated and publicly available, published data sets consisting of 17,629 gene names,^94, 95^ resulting in a mouse data set of 16,537 genes by 186,900 nuclei. Raw human and mouse data were extracted from processed objects for integration.

#### Optimization and Integration

To select lambda and k parameters, lambda optimization was run for k = 20, 25, 30, while k optimization was done with lambda set to 15, 20, and 30 (Fig. 1SA-B). For final analyses, the choice was lambda = 30 and k = 20, 25, 30. LIGER^14^ with R package rliger^96^ was used to integrate raw human and mouse data processed as described above, with 2,000 variable genes, lambda = 30, k = 20, 25, 30, with 3 restarts and maximum iteration of 100. After quantile normalization and UMAP (Uniform Manifold Approximation and Projection) dimension reduction for visualization (Fig. S1C), alignment and agreement were calculated, and the object was converted to Seurat for downstream analyses. Overall alignment and agreement metrics were: 0.7152 and 0.0199, 0.7557 and 0.0298, 0.7193 and 0.0324 for k = 20, 25, and 30, respectively. Integrated object with highest alignment metric, k=25, was chosen for downstream analyses (with individual cluster alignments in Table S1).

### Integrated dataset analyses

Cell type identities for each cluster in the integrated data set were assigned using a combination of R package MetaNeighbor^97^ and differential gene expression analysis.^13^ Reference data set previously trained on the Brain Initiative Cell Census Network (BICCN) mouse primary motor cortex data sets was utilized in MetaNeighbor as previously described^98^ to determine best cell type match for each cluster. The AUROC (area under the receiver operator characteristic curve) was calculated for each cluster, with an average of 0.86 ± 0.10 for highest AUROC (Table S2). Neuronal clusters were classified as excitatory or inhibitory based on SLC17A7/SLC17A6, and GAD1/GAD2 expression, respectively.

### Stratifying mouse populations based on memory performance

To define resilient and susceptible groups, transgenic mice at 14 months of age were ranked according to their individual CFM performance as reported previously^10^ (Fig. 1B). Mice that performed below average were classified as susceptible, and mice that performed above average were classified as resilient (Fig. 1B).

### Differential gene expression analyses

Differential gene expression analysis was performed on human and mouse samples separately. For analysis of resilience signatures, genes that were expressed in at least 10% of nuclei in least one group (resilient or susceptible) were analyzed in the data from 14-month-old transgenic mice and human subjects with high pathology. Differential expression statistics for each cluster were performed with glmmTMB function^99^ that uses generalized linear mixed model (GLMM) using Template Model Builder (TMB) with family function nbinom2(link = “log”) with cognition status (resilient or susceptible) as fixed effect and with each individual as the random effect, and ANOVA function in the car package.^100^ Multiple testing adjustment was calculated for each cluster with the Benjamini–Hochberg procedure using p.adjust function of the stats package in R.^91^ Upregulated and downregulated genes were those whose log_2_FC is equal to or greater than 0.25, or equal to or less than −0.25, respectively. Gene ontology (GO)^101, 102^ enrichment analysis on upregulated or downregulated genes was performed with clusterProfiler package with all gene set sizes and against all background genes for broad categories, and using gene sizes of under 500 for sub-classification.^103^

For analysis of resilience genes, the same generalized linear mixed model analysis as above was performed on pre-selected candidate genes in four groups: 6-month-old non-transgenic mice, 14-month-old non-transgenic mice, 6-month-old transgenic mice, and 14-month-old transgenic mice.

### Immunohistochemistry

The immunohistochemistry study incorporated a different cohort of male and female 14-month-old AD-BXD mice from the following strains: 14, 16, 22, 32, 44, 55, 56, 60, 61, 66, 68, 75, 77, 81, 87, 89, 99, 100, as well as the B6 and D2 F1s. For the female-only dataset, samples from the following AD-BXD strains were included: 14, 16, 22, 55, 75, 77, 81, 99, as well as the B6 and B6*D2 F1s. After CFM testing, mice were anesthetized with isoflurane and decapitated. The brains were removed and halved; one hemibrain was placed in 4% paraformaldehyde and kept at 4°C until the samples were sent to Neuroscience Associates (NSA, Knoxville, TN) for processing. The hemibrains were embedded, processed, and stained simultaneously in blocks of 40. The brains were freeze-sectioned coronally at 40 μm intervals (not including the cerebellum). Serial sections were stained for neurons using NeuN (anti-NeuN antibody, clone A60, biotin conjugated, Millipore Cat. No. MAB377B, 1500 dilution) and visualized using 3,3′-Diaminobenzidine (DAB), resulting in 22 images per brain on average. Images were taken with 20x objective on a Huron Digital Pathology TissueScope LE120 (0.4 microns/pixel).

### Image analysis

Cropped and down-sampled images from hemibrains of 29 mice were systematically registered to the Allen Brain Atlas CCFv2017^104^ and NeuN coverage across layers 4 and 5 of designated cortical regions were quantified using the QUINT workflow.^105–107^ Neuronal coverage in these regions was assessed as a measure of the area of stain coverage over the area of the region. Regions of the frontal cortex were identified and neuronal coverage within layers 4 and 5 were averaged from all sections per brain. Layers 4 and 5 from the following areas of the frontal cortex were assessed: prelimbic area, infralimbic area, anterior cingulate area (dorsal part), anterior cingulate area, anterior cingulate area (ventral part), agranular insular area (dorsal part), agranular insular area (ventral part), primary motor area, secondary motor area, primary somatosensory area, somatosensory areas, orbital area (lateral part), orbital area (medial part), dorsal peduncular area, and frontal pole.

Resilient and susceptible status for each strain was assigned using CFM averages across the male and female 14-month-old AD-BXD data set.^10^ Strains that fell below the population average were deemed susceptible while those scoring above the average were deemed resilient to cognitive decline. Cognitive status was then correlated with strain averages of neuronal coverage from each layer 4 and 5 frontal cortical region using a biserial correlation.

### Deconvolution

For deconvolution of bulk RNA-seq data to obtain estimates of cluster proportions, we used the dtangle R package.^17^ This analysis consisted of the following steps:

1. We created pseudobulk samples from the snRNA-seq data set, starting with nuclei having >500 genes detected. The counts from these nuclei were then added together in random combinations, such that each cluster was represented between 0-30% (in increments of 2%) in at least 10 pseudobulk samples.
2. We then ran dtangle on these pseudobulk samples, with the following combinations of parameters: n_markers in [0.01,0.02,0.03,0.05,0.1,0.2], marker_meth in [“ratio”, “diff”, “p.value”], and either with or without CPM normalization of the pseudobulk and single-nucleus RNA-seq counts. The reference set for dtangle was the full single-nucleus RNA-seq data set.
3. We selected an optimal set of parameters for dtangle based on which pseudobulk predictions had the highest correlation with the ground truth cluster proportions (used to generate the pseudobulk samples). The correlation for each cluster was calculated separately for each parameter set, and the parameter set yielding the highest minimum correlation value over all clusters was chosen. This resulted in n_markers=0.1, marker_meth=”ratio”, and CPM normalization.
4. We ran dtangle using these parameters on the bulk frontal cortex RNA-seq data set to obtain final estimates of the clusters. Bulk RNA-seq data for an AD-BXD mouse panel comprising of 235 mice of 28 different strains including male, female, 6 and 14 mo, age mice were used, with 18 samples filtered for this data set to match the snRNA-seq data set (female, 5XFAD, and 14 mo) for analysis. CFM data was used as described above to classify all strains in this study as resilient or susceptible, finally resulting in 10 resilient and 8 susceptible samples. Frontal cortex RNA-seq data and the corresponding behavioral data is deposited on SAGE (https://www.synapse.org/#!Synapse:syn17016211).

### Human references validations

To evaluate the relevance of resilience candidates to resilience in independent human cohort studies, our team leveraged published genomic, transcriptomic, and proteomic data. Both human and mouse genes were evaluated, with reported values for mouse genes included in this report. First, we leveraged data from a published genome-wide association study (GWAS) of resilience to AD neuropathology, defined as better-than-predicted cognitive performance given an individual’s amyloid burden.^78^ PrediXcan^108^ was used to quantify predicted levels of 61 candidate genes across 28 tissue types leveraging the GTEx database for model building and applied using GWAS data. Tissue-specific expression models were built leveraging elastic-net regression in the cis gene region (within 1Mb) and selected based on five-fold cross-validation as previously described. We then regressed our published resilience trait (n=5108) on each gene model covarying for age and sex. Correction for multiple comparisons was completed leveraging the false discovery rate (FDR) procedure (correcting for all gene-tissue combinations).

Next, we leveraged bulk transcriptomic data from the ROSMAP to evaluate whether the expression of resilience genes also related to cognitive performance in the years preceding death. ROSMAP enrolled older adults without dementia who agree to annual clinical evaluations and brain donation at death.^109^ Bulk RNA sequencing was performed in 3 brain regions: the head of the caudate nucleus (CN), dorsolateral prefrontal cortex (DLPFC), and posterior cingulate cortex (PCC), all of which were processed following a published protocol.^110^ A global cognitive composite was calculated by averaging z-scores from 17 tests, as previously described.^111^ We evaluated transcript associations with cross-sectional cognition covarying for age at death, post mortem interval, and sex, and we evaluated associations with longitudinal cognition leveraging mixed effects regression models with the same covariates and both the intercept and slope (years from death) entered as fix and random effects in the model. Correction for multiple comparisons was completed with FDR procedure.

Finally, to identify proteins that shared resilience signatures in the human ROSMAP and Banner data set, data from Johnson et al.^52^ was used (Table 6, considering p- and bicor values for MMSE30). Proteins that were upregulated in AsymAD vs AD can be found in Supplementary Table S2 of that publication. For both data sets, in cases where multiple values were reported per protein, the proteins with the lowest *p-*values were chosen.

### Druggability rankings

Druggability of nominated genes were assessed using a protein druggability dataset (https://www.synapse.org/#!Synapse:syn13363443) combined with Agora AD gene nomination (https://agora.adknowledgeportal.org/genes). R package biomaRt was used to match gene names with their ensembl IDs.^112, 113^

Small molecules targeting genes of interest were identified using https://www.genecards.org/ resource.

### Identification of drug repositioning candidates

To identify drugs that might enable boosting/mimicking a gene expression signature linked to resilience mechanisms, we used several drug signature search algorithms as implemented in the signatureSearch package^114^ (v.1.12.0) in R-4.2.2. As reference databases for signature searches, we used both the Connectivity MAP (CMAP) database^115^ and the Library of Integrated Network-based Cellular Signatures (LINCS)^116^ 2020 L1000 dataset. We employed four search methods for CMAP (the CMAP search method, the LINCS search method, the gCMAP search method, and correlation-based) and three for LINCS2020 (gCMAP did not yield meaningful results). We used default parameters for all methods except for gCMAP, where we set higher=0, lower=NULL, and padj=0.05 to limit the search space to (significantly) upregulated gene signatures.

We then first conducted ensemble rank aggregation of results for the different search strategies separately for CMAP and LINCS. To this end, we first extracted the most significant hit for each compound across screens and separated results into significant and non-significant signature matches as follows:

1. for the CMAP search method, we considered results as significant, if the Kolmogorov-Smirnov statistic was surpassing the critical value for P = 0.05 for distributions corresponding to the overlap of 52 (CMAP) and 53 (LINCS2020), respectively, genes;
2. for the LINCS search, we performed a post-hoc FDR correction after selecting the best result for each compound across screens and considered results with an FDR-corrected P ≤ 0.05 significant;
3. for gCMAP, we considered results as significant that surpassed an effect value corresponding to the smallest scaled Kolmogorov-Smirnov statistic still significant in the CMAP search, thus homogenizing results between the two search methods;
4. for the correlation-based signature search, we calculated the t-statistic from the correlation coefficient and considered results significant that had a two-tailed P ≤ 0.05.

Significant compounds were ranked according to their effect size in mimicking the resilience signature, while insignificant compounds were set to share the highest rank. Ranks were then combined using the Dowdall rule, an alternative approach to the Borda Count, where the top rank is transformed to 1/1, the second rank to 1/2, and the nth rank to 1/n. We then summed these scores separately for CMAP and LINCS, scaled the resulting scores for each to [0,1]- intervals and summed them up to retrieve the final overall ranking score. All compounds that had an overall ranking score ≥ 0.1 were considered overall significant.

Finally, we performed drug set enrichment (signatureSearch function dsea_hyperG) analysis for the 55 top compounds having target gene annotations available using the three ontologies (biological process, molecular function, and cellular component) of the Gene Ontology^101, 102^ (ExperimentHub version 2.6.0, dataset EH3232, added on 2019/10/22) using the Benjamini-Hochberg FDR-correction procedure to account for multiple testing.

### Statistical analysis and software

For Fig. 2A, R package corrplot^117^ was used on log of average expression + 1. Correlation values for Fig. 2B were calculated and graphed with stat_poly_eq() function from R package ggpmisc^118^ on genes with expression in at least one of the groups.

For Figs. 3A and B and S2D, cluster frequencies (Fig. 3) or cluster proportions (Fig. S2D) were calculated with all groups; then, R package rstatix^119^ was used to compare resilient and susceptible groups, t-test was performed for each cluster, with Benjamini-Hochberg multiple comparison correction.

For Fig. S2C, cognitive status of strains based on strain average CFM (resilient or suspectable) was correlated with strain averages of neuronal coverage from each of the layers 4 and 5 in the frontal brain region using a biserial correlation. Correlations between regions for the heatmap were run using rcorr from Hmisc in R.^120^ The correlation between regions and the binary R vs S status was completed using a cor.test (method = pearson) in R.

## Supplemental Information: Figure Legends

**Fig. S1. Parameter optimization for LIGER integration. A**. KL divergence plot varying lambda (15, 20, 30) and **B.** alignment plot varying k (20, 25, 30) for parameter selection for LIGER integration; **C.** Integrated human and mouse data set with lambda = 30, varying k (20 (left), 25 (middle), 30 (right)): cluster proportion by species, UMAP visualization of the integrated data set with cluster numbers, and separate UMAP of the data for mouse (top) and human (bottom).

**Fig. S2**. **A.** Log of nuclei numbers in each cluster for each sample; **B.** Dotplot showing expression of pan-neuronal, neuronal subtype, and layer-specific marker genes for neuronal clusters in the integrated data set; **C.** Correlation plot for NeuN coverage among different layer 4 and 5 regions and cognitive status in AD-BXD strains; **D.** No differences in cluster proportion estimates from bulk RNA-seq data between resilient and susceptible groups in AD-BXDs.

**Fig. S3. A.** Companion plot for Fig. 5 for each gene membership in top 10 GO:BP pathways. **B.** Expression level for neurogenesis-related genes in excitatory neuronal cluster 20 and inhibitory neuronal cluster 5.

**Fig. S4.** Gene membership in top 20 GO:BP pathways among **A.** nervous development genes from Figs. 5 and S3A and **B.** transport genes from Figs. 5 and S3A.

**Fig. S5.** Gene membership in top 20 GO:BP pathways among downregulated genes in **A.** C1 endothelial human cluster for resilient group compared to susceptible group and **B.** C11 immune – other mouse cluster for resilient group compared to susceptible group.

