## Supplementary Figures for "Conserved cell-type specific signature of resilience to Alzheimer’s disease nominates role for excitatory intratelencephalic cortical neurons"

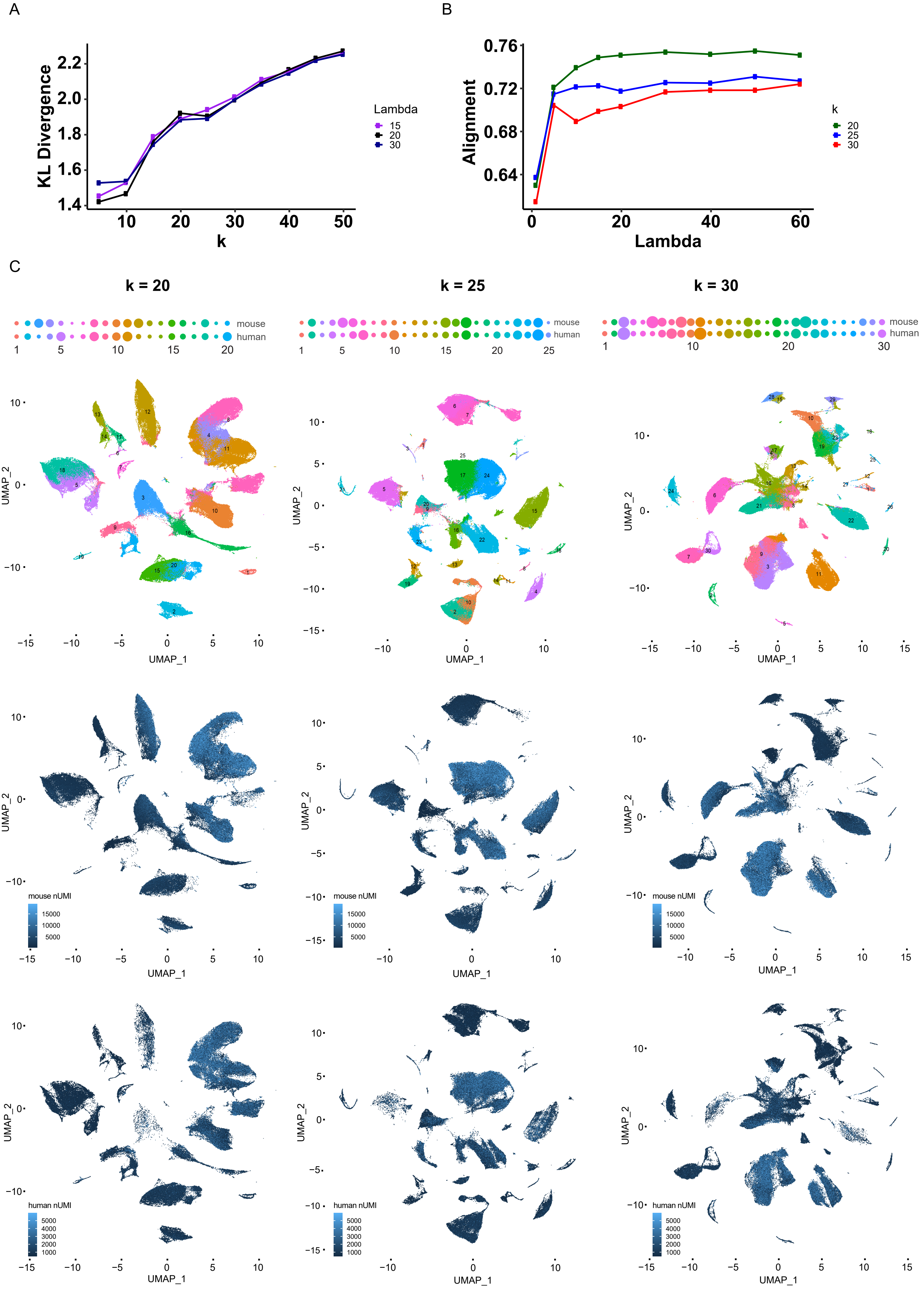


Supplementary Figure 1

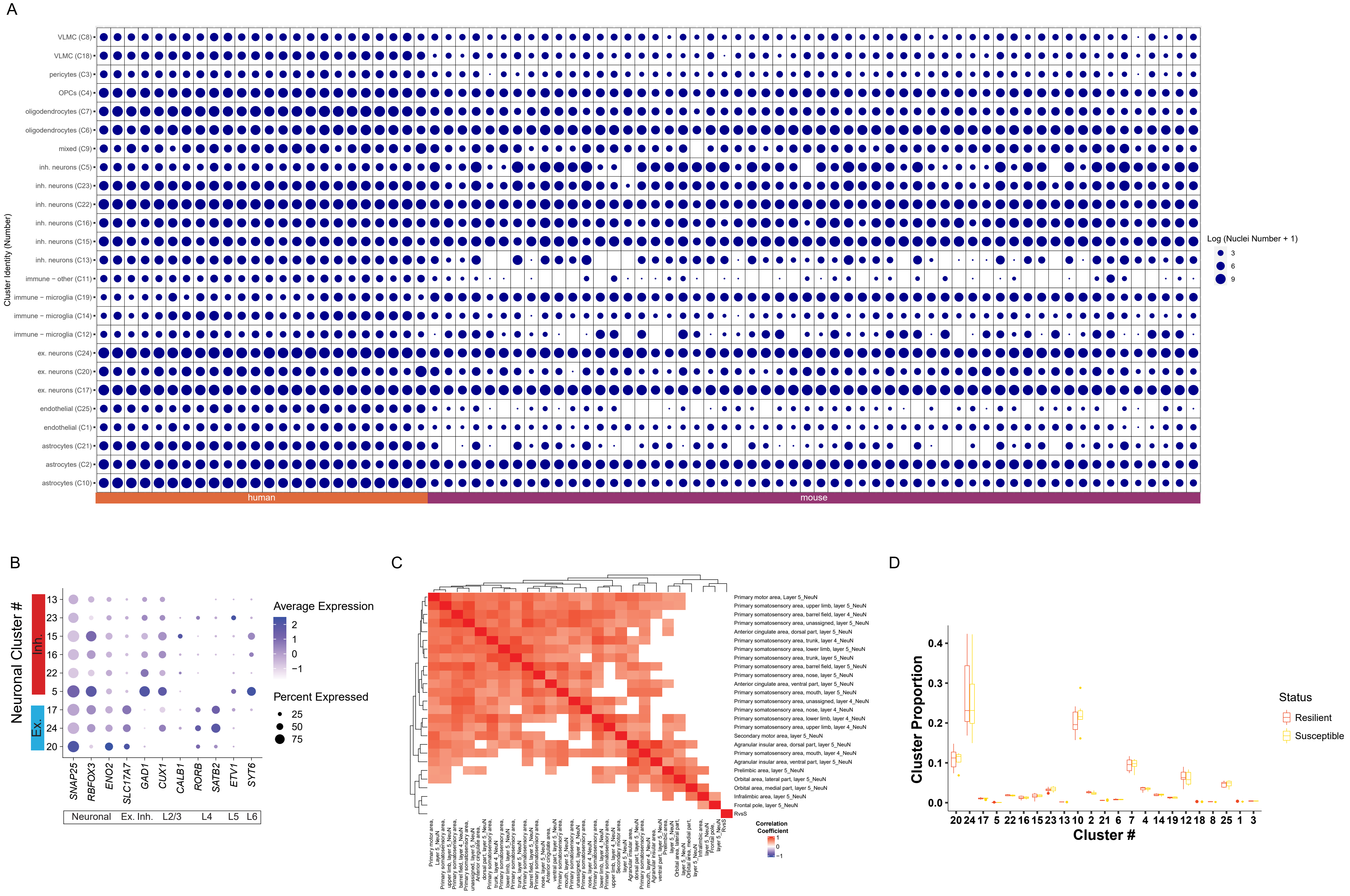


Supplementary Figure 2

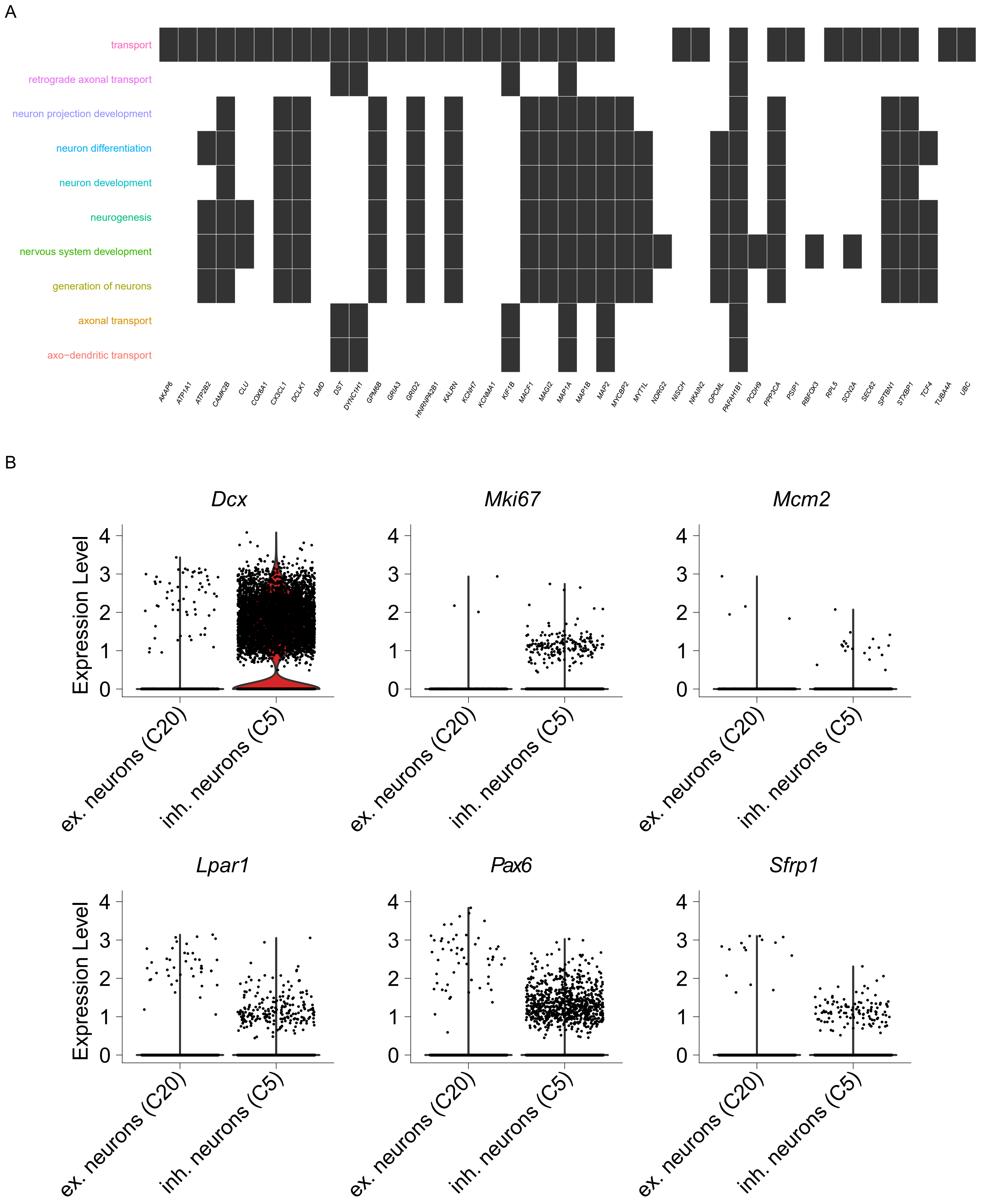


Supplementary Figure 3

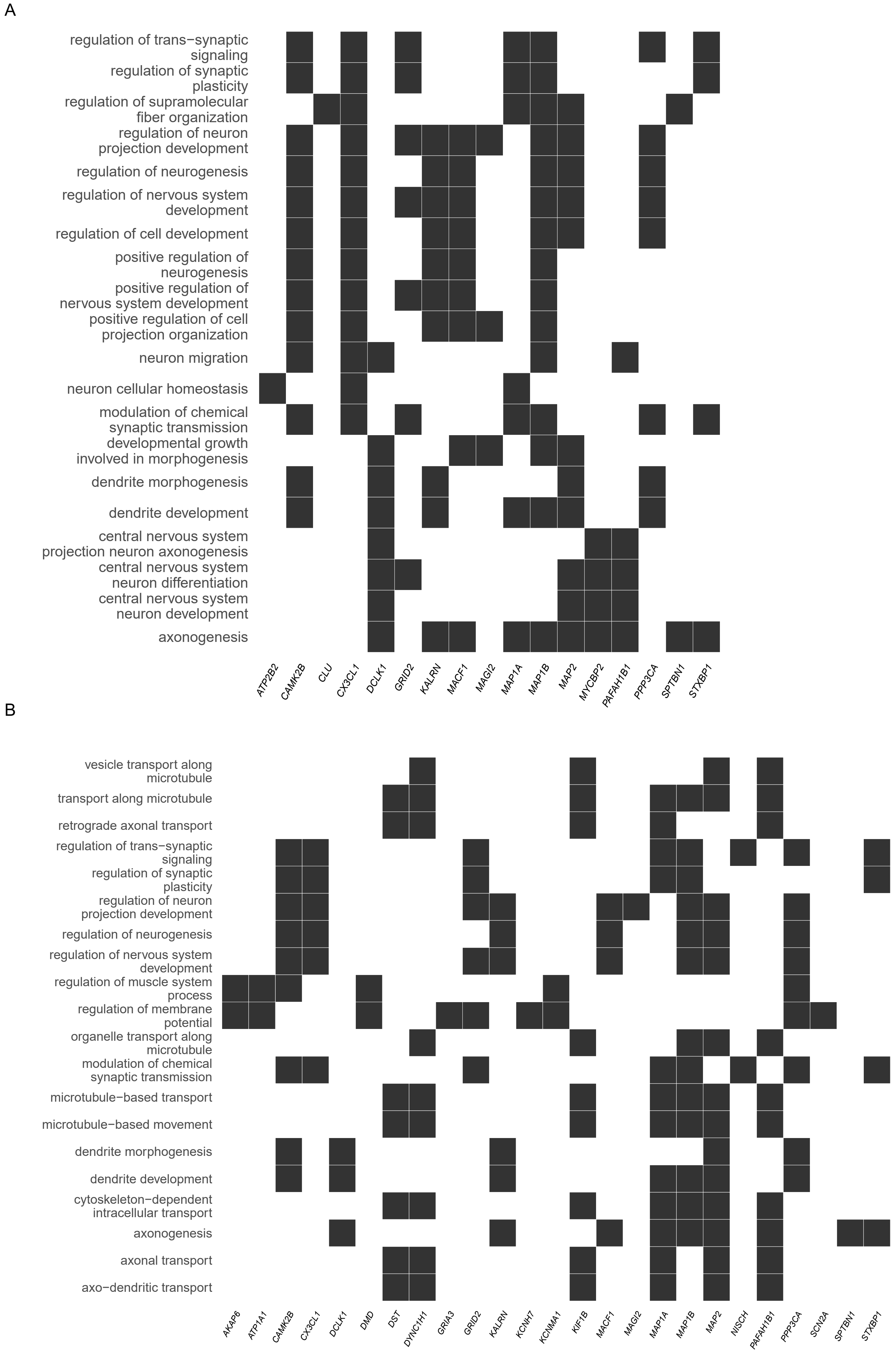


Supplementary Figure 4

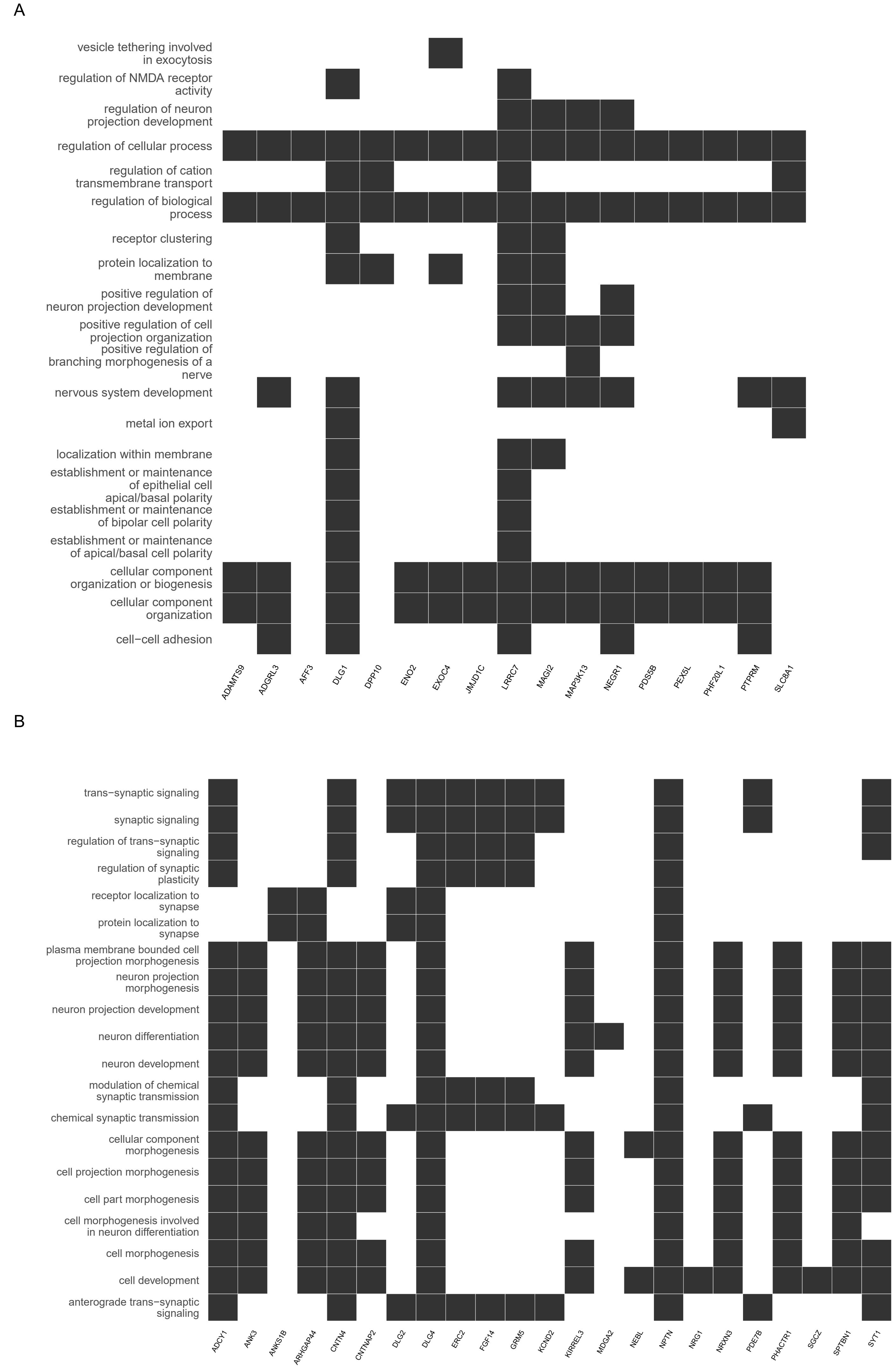


Supplementary Figure 5
